# Regional Differences in Dorsal Skin Determine the Rate of Adhesive Material-Induced Hair Regeneration

**DOI:** 10.64898/2026.07.22.739732

**Authors:** Tomoyuki Toge, Moyuri Inoue, Ying-Ying Chou, Anna Yoshimura, Sachiko Yamashita Takeuchi, Kayoko N. Kuroishi, Kaori K. Gungikake, Jun J. Miyamoto, Takuma Matsubara, Osamu Kaminuma, Tatsuo Kawamoto, Shoichiro Kokabu

**Affiliations:** Division of Biochemistry, Kyushu Dental University, Kitakyushu, Fukuoka 803-8580, Japan; Division of Orofacial Functions and Orthodontics, Faculty of Dentistry, Kyushu Dental University, Kitakyushu, Fukuoka 803-8580, Japan; Division of Applied Pharmacology, Faculty of Dentistry, Kyushu Dental University, Kokurakita-ku, Kitakyushu 803-8580, Japan; Department of Disease Model, Research Institute of Radiation Biology and Medicine, Hiroshima University, Hiroshima 734-8551, Japan

**Keywords:** Adhesive material, cyanoacrylate, dorsal skin, hair cycle, Hox genes, regional heterogeneity

## Abstract

Dorsal skin is widely used in mouse wound-healing, dermatitis, and hair-cycle models, but is often treated as a uniform site. We investigated whether adhesive material (cyanoacrylate)-induced hair regeneration, a model we previously reported, differs by position within the dorsal skin. Adhesive material was applied to four regions along the cranial-to-caudal axis of the mouse dorsal skin. Hair regrowth appeared earlier at the cranial sites than at the caudal sites, and this difference persisted through late anagen and the anagen-to-catagen transition. In contrast, hair regrowth after full-thickness skin excision was slower, smaller in area, and less reproducible than that after adhesive material application. Expression of Hox genes, including Hoxa9, Hoxb9, Hoxc9, Hoxa10, and Hoxc10, was higher at the caudal sites than at the cranial sites but did not correlate with the timing of hair regrowth. RNA sequencing of intact skin from the cranial and caudal sites revealed distinct baseline profiles, including differences in the Wnt inhibitor Sfrp4 and the adipogenic genes Pparγ and Fabp4. Early after adhesive material application, histological changes and the expression of inflammation- and tissue-repair-related genes also differed between the cranial and caudal skin. These findings indicate that mouse dorsal skin is not a uniform experimental field and that cranial-to-caudal position should be considered when designing and interpreting hair-regeneration and wound-healing experiments in mice.

## Introduction

Dorsal skin is used in a wide range of experimental systems, including wound healing, dermatitis, skin barrier function, evaluation of topical agents, and hair growth/hair-cycle research [4,12,19]. The dorsal region is also widely used for subcutaneous tumor transplantation and for the analysis of tumor angiogenesis and microcirculation using dorsal skinfold chamber models [14,15]. At the subcutaneous level, the dorsal region is also used for fibrosis models and for the analysis of adipose and mesenchymal tissues [20].

Many of these studies treat dorsal skin collectively as “dorsal skin” or “back skin.” However, dorsal skin is not necessarily uniform. Regional histological and functional differences in mouse skin have been reported previously. For example, anatomical sites such as the ear, tail, footpad, and back differ in epidermal and dermal structure, stratum corneum, the presence and density of hair follicles, and immune cell distribution [18]. Differences in coat color, hair length, hair type, and skin development along the dorsoventral axis have also been reported, indicating that skin has region-specific characteristics according to anatomical position [1]. In hair-cycle research, the hair cycle in adult mouse dorsal skin has been reported to progress as spatial hair-cycle domains and regenerative hair waves [11]. Plikus et al. further showed that dermal Bmp2/Bmp4 expression fluctuates periodically, generating functionally distinct states — a BMP-high refractory telogen and a BMP-low competent telogen — and that this BMP signal fluctuates in antiphase with Wnt/β-catenin signaling to regulate the activation state of hair follicle stem cells [12]. These findings indicate that dorsal skin is not a single uniform experimental field, but a tissue whose local environment — comprising the hair follicles, dermis, and subcutaneous adipose layer — varies in space and time.

Hox genes are a family of transcription factors that specify positional information along the body axis during development and are major determinants of anatomical regional identity in vertebrates [9]. HOX gene expression in adult skin also differs by anatomical site and has been reported to contribute to the positional memory of dermal fibroblasts and to site-specific epithelial responses [13]. Region-specific Hox-related effects have also been reported in hair follicles: Yu et al. showed that Hoxc-dependent mesenchymal niche heterogeneity of dermal papilla cells alters the regenerative activity of hair follicle epithelial stem cells and contributes to site-specific hair growth capacity [22]. These findings suggest that Hox genes may serve as a molecular indicator of regional identity and follicular responsiveness when examining differences in hair growth response along the cranial-to-caudal axis of dorsal skin.

We recently reported a new experimental model in which local application of an adhesive material induces hair growth in mouse skin [8]. In that model, localized hair induction was observed not only in dorsal skin but also in the head, thigh, and abdominal skin. However, whether regional differences within the dorsal skin itself affect the pattern of hair growth or the histological response to this method had not been examined. It was also necessary to compare hair induction by this method with that induced by an existing model of small full-thickness skin excision wounds, which is used to evaluate the telogen-to-anagen conversion triggered by wound stimulation [6].

In this study, we therefore focused on regional differences within mouse dorsal skin and examined whether the responsiveness of the local adhesive-material-induced hair-regeneration method differs by position within the dorsal skin. We further analyzed the telogen-to-anagen conversion induced by a small full-thickness skin excision wound, a representative existing hair-induction system, and compared it with adhesive-material-induced hair induction. In addition, we analyzed baseline gene-expression profiles in cranial and caudal dorsal skin and the early tissue response after adhesive material application.

## Materials and Methods

### Animals

Eight-week-old male and female C3H/HeSlc mice were purchased from Japan SLC, Inc. (Hamamatsu, Japan) and used in this study. Mice were maintained under standard laboratory conditions with free access to food and water. All animal experiments were approved by the Experimental Animal Care and Use Committee of Kyushu Dental University (approval number 26-004) and were conducted in accordance with the relevant institutional guidelines and regulations. All animal studies complied with the ARRIVE guidelines.

### Adhesive material-induced hair regrowth model and regional analysis of dorsal skin

Cyanoacrylates (adhesive materials) were obtained from Toagosei Company, Limited (Tokyo, Japan) and used as supplied by the manufacturer. Mice were anesthetized under general anesthesia using a mixture of medetomidine (0.75 mg/kg), midazolam (4 mg/kg), and butorphanol (5 mg/kg), as previously described [16]. The dorsal hair was shaved using clippers, and adhesive material was applied to defined areas of the dorsal skin. The day of adhesive material application was defined as Day 0. Macroscopic images of the dorsal skin were obtained under general anesthesia at the indicated time points after application. For regional analysis, the dorsal skin was divided into four sites along the cranial-to-caudal axis, designated Site 1, Site 2, Site 3, and Site 4. Site 1 represented the most cranial region, and Site 4 represented the most caudal region. Adhesive material was applied to each site, and the timing and extent of hair regrowth were compared among the four sites.

### Full-thickness skin excision assay

To examine regional differences in the gross wound-healing process and subsequent hair regrowth after skin injury, and to compare this response directly with adhesive material-induced hair regrowth within the same animal, circular full-thickness skin excision wounds were generated at the Site 1- and Site 4-equivalent positions on one side of the dorsal skin under general anesthesia [6], while adhesive material was applied to the corresponding cranial-to-caudal positions on the contralateral side of the same animal. A 6-mm biopsy punch (KAI Corporation, Tokyo, Japan) was used to mark and initiate circular full-thickness skin excisions, and scissors were used to complete the excision when necessary. The external appearance of wound closure and subsequent hair regrowth was documented by serial gross photography at the indicated time points.

### Histopathological examination

Skin samples were collected from the indicated dorsal skin sites and fixed with 4% paraformaldehyde (FUJIFILM Wako Pure Chemical Corporation, Osaka, Japan) in PBS. Samples were dehydrated through an ethanol and xylene series, embedded in paraffin, and cut into 4-μm sections. After deparaffinization, sections were stained with hematoxylin and eosin. Histological images were acquired using a Keyence BZ-X800 microscope (Keyence) [21].

### Quantitative real-time PCR

Total RNA was isolated from skin tissues using a FastGene RNA Basic Kit (Nippon Genetics, Tokyo, Japan) and reverse-transcribed into cDNA using a High-Capacity cDNA Reverse Transcription Kit (Applied Biosystems, Waltham, MA, USA). SYBR Green-based quantitative real-time PCR was performed using PowerUp SYBR (Thermo Fisher Scientific, Waltham, MA, USA) and the QuantStudio 3 Real-Time PCR System (Thermo Fisher Scientific) [7]. Relative gene expression was calculated by the ΔCT method using *Tbp* as the housekeeping gene for normalization. For regional Hox gene expression analysis, skin samples were collected from Site 1, Site 2, Site 3, and Site 4 of 5 animals (n = 5 biologically independent animals), and the expression levels of *Hoxa9*, *Hoxb9*, *Hoxc9*, *Hoxa10*, and *Hoxc10* were analyzed; each sample was measured in technical triplicate by qPCR (n = 3 technical replicates per animal). For analysis of adhesive material-induced molecular responses, skin samples were collected from Site 1 and Site 4 of 5 animals (n = 5) at the indicated time points, and the expression levels of *Sfrp4*, *Pparγ*, *Fabp4*, *Nos2*, *Tnfα*, *Cd86*, *Arg1*, *Igf1*, and *Mrc1* were analyzed, each in technical triplicate (n = 3). Primer sequences are listed in Supplementary Table 1.

### RNA sequencing and bioinformatics analysis

Total RNA was extracted from paired Site 1 and Site 4 dorsal skin samples collected from the same 2 animals (n = 2 animals total; both sites were sampled from each animal, yielding 2 samples per site). RNA sequencing and initial bioinformatic analysis were performed by Novogene Co., Ltd. Messenger RNA was purified from total RNA using poly-T oligo-attached magnetic beads. After fragmentation, first-strand cDNA was synthesized using random hexamer primers, followed by second-strand cDNA synthesis for strand-specific library construction. The libraries were subjected to end repair, A-tailing, adapter ligation, size selection, amplification, and purification. Library quality was assessed using Qubit, real-time PCR, and a bioanalyzer, and the quantified libraries were sequenced on an Illumina platform. Raw reads in FASTQ format were subjected to quality control. Reads containing adapter contamination, reads containing more than 10% uncertain nucleotides, and low-quality reads were removed to obtain clean reads. Sequencing quality was evaluated using the error rate, Q20, Q30, and GC content. Clean reads were aligned to the mouse reference genome mm39 using HISAT2. Gene expression levels were quantified using read counts and FPKM values. Differential gene expression analysis between Site 1 and Site 4 was performed using DESeq2. Differentially expressed genes were identified using thresholds of adjusted P value ≤ 0.05 and |log2FoldChange| ≥ 1. Gene Ontology and KEGG pathway enrichment analyses were performed using clusterProfiler to identify biological processes and pathways associated with region-specific gene expression profiles.

### Statistical analysis

Data are presented as the mean ± SD. Comparisons between two groups were performed using an unpaired two-tailed Student’s t-test. Comparisons among multiple groups were performed using one-way ANOVA followed by Tukey’s multiple-comparison test. In Figure 3, Hox gene expression at Site 2, Site 3, and Site 4 was compared with Site 1. In Figure 5, gene expression at Site 4 was compared with Site 1 at the same time point. *P < 0.05 and **P < 0.01 were considered statistically significant.

## Results

### Adhesive material-induced hair regrowth differs along the cranial-to-caudal axis of dorsal skin

We first examined whether adhesive-induced hair regrowth shows regional differences. The dorsal skin of mice was divided into four regions from the cranial to the caudal side, designated Site 1, Site 2, Site 3, and Site 4, and adhesive material was applied to each site (Figure 1A, 1B). In both male and female mice, localized hair regrowth was induced at the sites of adhesive material application, and this response did not proceed uniformly across the entire dorsal skin: it appeared earlier at the cranial sites, Site 1 and Site 2, and later at the caudal sites, Site 3 and Site 4. No clear sex difference was observed in this regional pattern, and male mice were used in all subsequent experiments.

**Figure 1.**
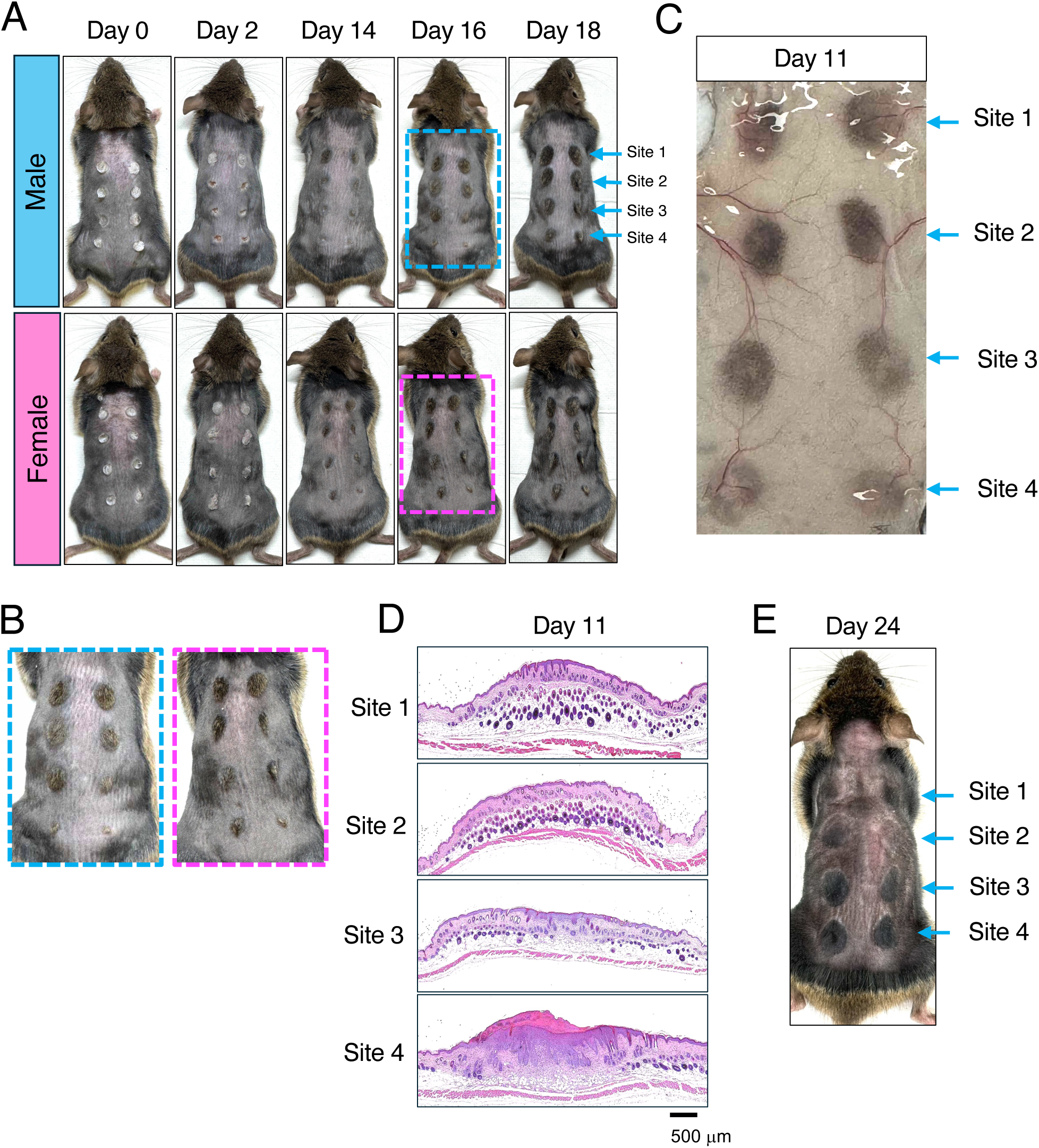
Adhesive material-induced hair regrowth is observed earlier in cranial than in caudal dorsal skin. The dorsal skin of 8-week-old male and female mice was divided into four regions from the cranial to the caudal side (Site 1-4), adhesive material was applied to each site, and the hair-regrowth response was observed over time. (A) Representative gross images of male and female mice after adhesive material application. Hair regrowth was observed earlier at Site 1 and Site 2, corresponding to the cranial dorsal skin, than at Site 3 and Site 4, corresponding to the caudal dorsal skin. (B) Enlarged images of the areas outlined by the dashed boxes in (A), for the male and female mouse, respectively. (C) Dermal-side view of the dorsal skin on Day 11 after adhesive material application. (D) Hematoxylin and eosin-stained sections of skin from each site on Day 11 after adhesive material application. Scale bar: 500 μm. (E) Representative gross images of each site on Day 24 after adhesive material application. n = 5. All photographs show representative examples.

This regional difference in the hair regrowth pattern was more clearly observed when the dorsal skin was examined from the dermal side on Day 11 (Figure 1C). At the Day 16 gross appearance, clear hair regrowth was also observed at Site 1 and Site 2, whereas hair regrowth was relatively delayed at Site 3 and Site 4 (Figure 1B). Furthermore, histological examination of hematoxylin and eosin-stained sections showed marked hair follicle elongation at Site 1 and Site 2 on Day 11, whereas this was less pronounced at Site 3 and Site 4, confirming histologically that hair-cycle progression preceded in the cranial region (Figure 1D).

By Day 24, the hair cycle at Site 1 and Site 2 had progressed further, with an appearance suggesting transition from late anagen to catagen. In contrast, at Site 3 and Site 4, the delayed hair regrowth was still clearly observed (Figure 1E). Thus, the cranial-to-caudal time difference observed in the hair regrowth response after adhesive material application was not limited to a transient difference at the onset of hair regrowth, but was maintained through the late stages of hair-cycle progression. This suggests that the difference between Site 1/Site 2 and Site 3/Site 4 is less likely to reflect a large difference in the duration of anagen itself, and more likely reflects a difference in the timing of onset of the telogen-to-anagen transition, which is then reflected throughout the subsequent hair-cycle progression.

As with adhesive material-induced hair regrowth, we next examined whether the hair-regrowth response after full-thickness skin excision, another system for evaluating the telogen-to-anagen transition of resting hair follicles, also differs by region. After full-thickness skin excision, the wound decreased in size over time, and macroscopically apparent hair regrowth was observed around Day 18 after wounding; however, hair regrowth was absent at some sites, and stable hair regrowth was not obtained (Figure 2A). When directly compared with adhesive material-induced hair regrowth within the same animal, hair regrowth after full-thickness skin excision was confirmed later, was smaller in area, and included regions with no hair regrowth at all, compared with adhesive material-induced hair regrowth (Figure 2B). Wounds created at the Site 1-equivalent position tended to show earlier hair regrowth than those at the Site 4-equivalent position, but this difference was not reproducible enough to be evaluated with confidence. Hair-regrowth area and timing were also highly variable, and clear, localized hair induction was not observed (Figure 2B). Even at later observation points, hair regrowth was present in some cases and not clearly apparent in others, and the response remained unstable (Figure 2B).

**Figure 2.**
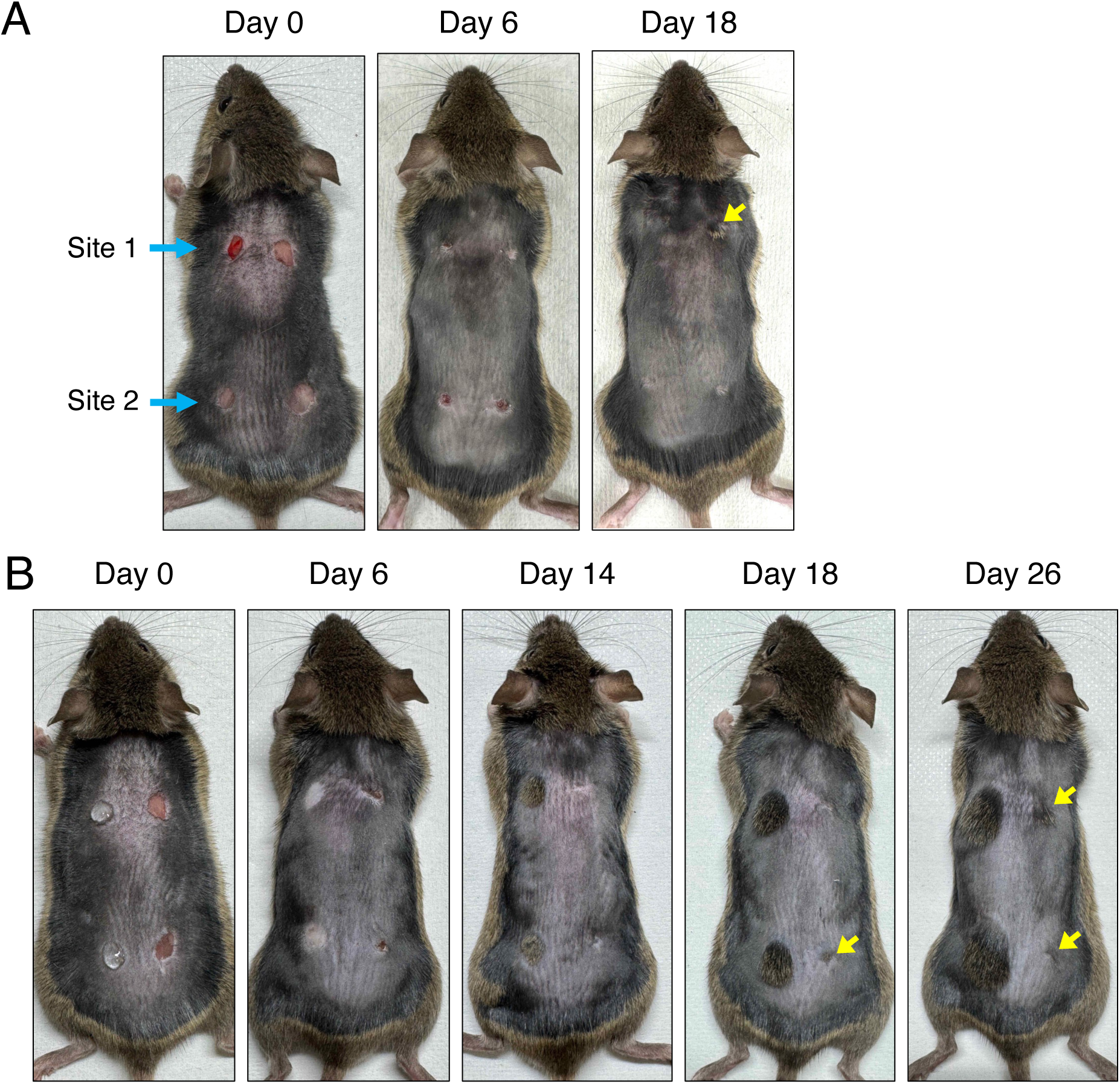
Wound healing and hair regrowth after small full-thickness skin excision are slower and less reliable than at adhesive material-treated sites. (A) Representative gross images after creating small full-thickness skin excision wounds at four sites. Day 0 shows the wounds immediately after creation; subsequent panels show the course of wound healing and hair regrowth on Day 6 and Day 18. Yellow arrowheads indicate sites of hair regrowth. (B) Adhesive material was applied to the left side of the dorsal skin, and small full-thickness skin excision wounds were created at the Site 1- and Site 4-equivalent cranial-to-caudal positions on the right side of the dorsal skin of the same animal. Representative images show the course of wound healing and hair regrowth on Day 0, 6, 14, 18, and 26 after treatment. Yellow arrowheads indicate sites of hair regrowth. n = 5. All photographs show representative examples.

### Cranial and caudal dorsal skin differ in tissue response and gene-expression profile

Next, to investigate why these cranial-to-caudal differences among Site 1 to Site 4 arise (Figure 3A), we examined the expression of the Hox genes *Hoxa9*, *Hoxb9*, *Hoxc9*, *Hoxa10*, and *Hoxc10* (Figure 3B). Most of the Hox genes examined, including *Hoxc10*, which showed the largest fold change, were more highly expressed at the caudal sites (Site 3 and Site 4) than at the cranial sites (Site 1 and Site 2) (Figure 3C). However, expression of these Hox genes was not higher at Site 1 and Site 2, where the adhesive material-induced hair regrowth response was early; rather, it was higher at Site 3 and Site 4, where the response was delayed. Thus, the expression pattern of the Hox genes examined in this study did not simply explain the timing of the adhesive material-induced hair regrowth response.

**Figure 3.**
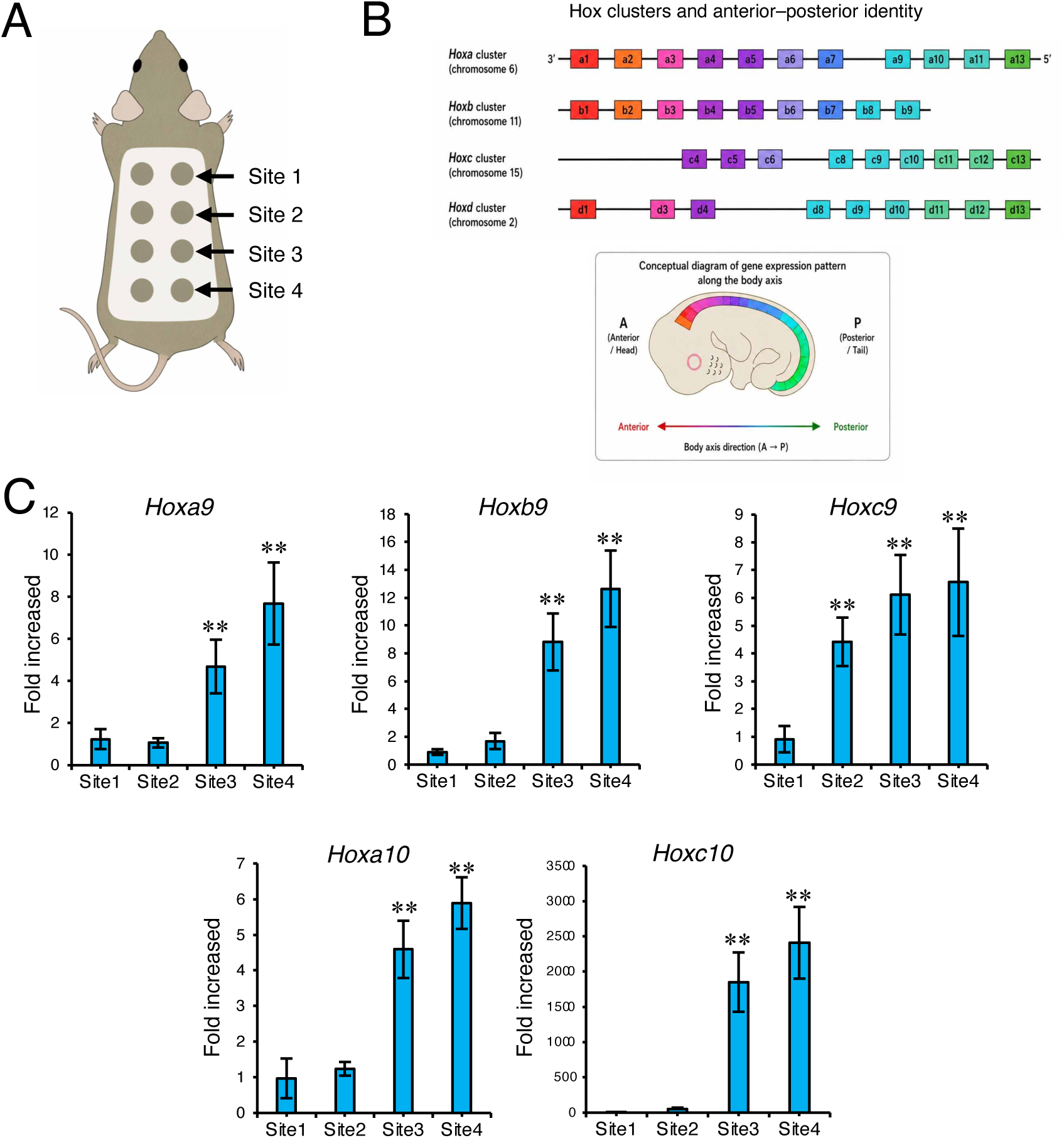
Hox gene expression changes along the cranial-to-caudal axis of dorsal skin. (A) Schematic diagram showing the positions of Site 1-4 in dorsal skin. (B) Schematic diagram showing the relationship between the Hox gene clusters and positional information along the anterior-posterior axis. (C) Expression analysis of *Hoxa9*, *Hoxb9*, *Hoxc9*, *Hoxa10*, and *Hoxc10* at Site 1-4. Several Hox genes showed distinct expression patterns from the cranial to the caudal dorsal skin. Data are presented as mean ± SD. **P < 0.01 vs. Site 1 (one-way ANOVA followed by Tukey’s multiple-comparison test). n = 5 animals; qPCR was performed in technical triplicate.

To investigate this further, we performed RNA sequencing using intact Site 1 and Site 4 skin (Figure 4A). Comprehensive comparison of gene expression between Site 1 and Site 4 identified multiple differentially expressed genes, indicating that the two sites have distinct gene-expression profiles (Figure 4B). Gene Ontology enrichment analysis using the differentially expressed genes extracted genes related to tissue morphogenesis, receptor ligand activity, and signaling receptor regulator activity, among others associated with skin tissue properties and tissue responses (Figure 4C). In addition, genes related to the adipose-metabolic niche showed distinct expression patterns between Site 1 and Site 4 (Figure 4D). qPCR analysis also showed altered expression of the Wnt-signaling inhibitor *Sfrp4* and the adipocyte differentiation-related genes *Pparγ* and *Fabp4* (Figure 4E). Analysis of GO terms and KEGG pathways related to inflammatory and wound response also identified pathways related to leukocyte migration, the IL-17 signaling pathway, and other inflammation-, metabolism-, and tissue-response-related processes (Figure 4F).

**Figure 4.**
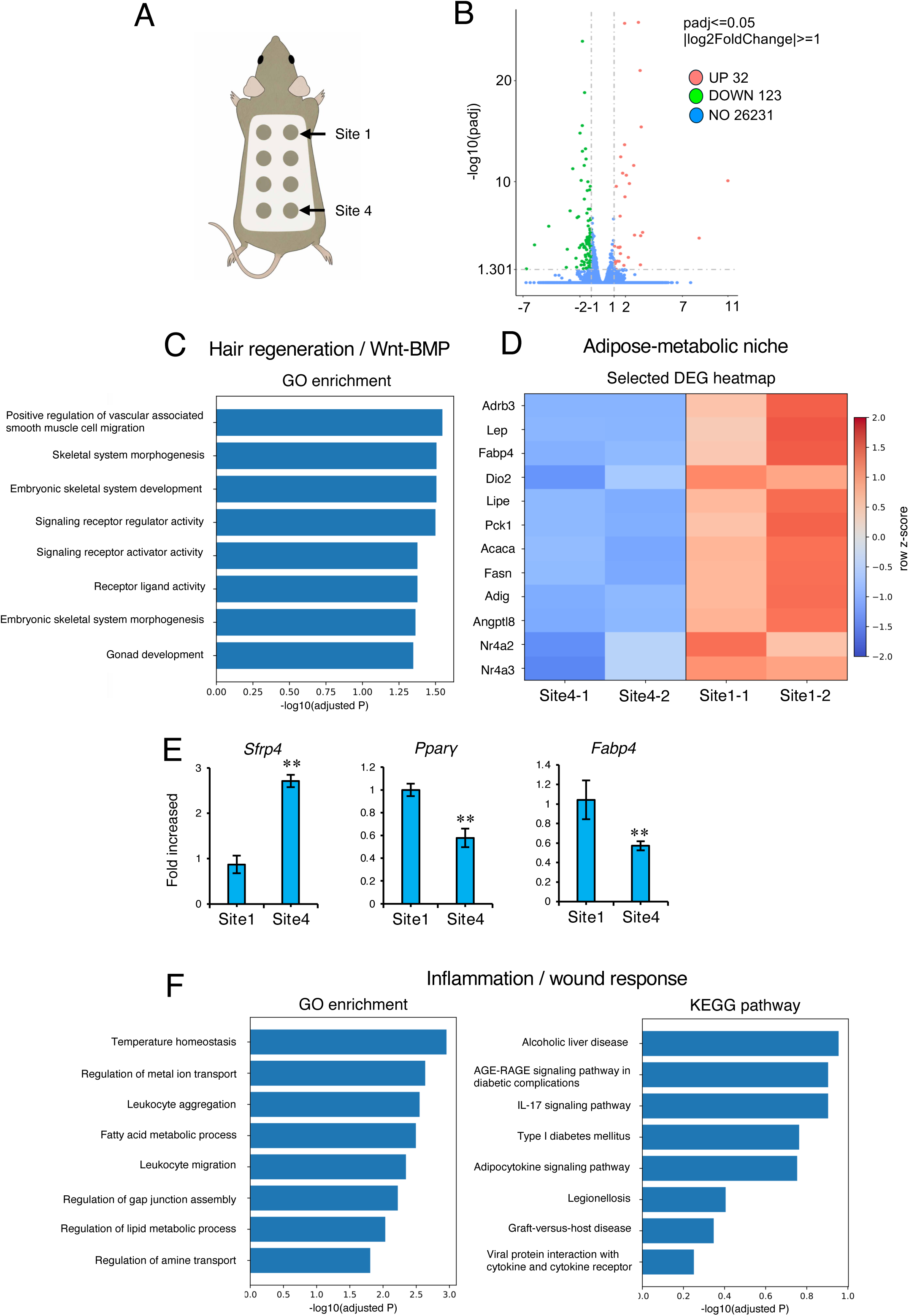
Cranial and caudal dorsal skin show distinct gene-expression profiles. RNA sequencing was performed using paired Site 1 and Site 4 skin samples from the same 2 animals (n = 2 animals total). (A) Schematic diagram showing the positions of the cranial (Site 1) and caudal (Site 4) dorsal skin used for RNA-seq analysis. (B) Volcano plot showing differences in gene expression between Site 1 and Site 4; upregulated, downregulated, and non-significantly changed genes were classified based on adjusted P value and log2 fold change. (C) Gene Ontology enrichment analysis, including genes related to hair regeneration, Wnt/BMP signaling, and skeletal formation. (D) Heatmap showing expression of genes related to the adipose-metabolic niche. (E) Expression analysis of *Sfrp4*, *Pparγ*, and *Fabp4* at Site 1 and Site 4. (F) Gene Ontology enrichment analysis and KEGG pathway analysis related to the inflammatory and wound response.

Because genes related to inflammation, metabolism, and tissue response were altered, we finally observed changes immediately after and shortly following adhesive material application. At Day 0, both Site 1 and Site 4 showed a telogen-like skin structure. After adhesive material application, tissue changes accompanied by epidermal thickening and inflammatory cell infiltration were observed from Day 1 to Day 2, followed by changes in hair follicle structure from Day 4 to Day 6 (Figure 5A). These tissue changes differed between Site 1 and Site 4: at the cranial site, where the hair-regrowth response was early, inflammation after adhesive material application resolved earlier, and hair follicle structure formed earlier. At the caudal site, where the hair-regrowth response was delayed, these changes were correspondingly delayed.

**Figure 5.**
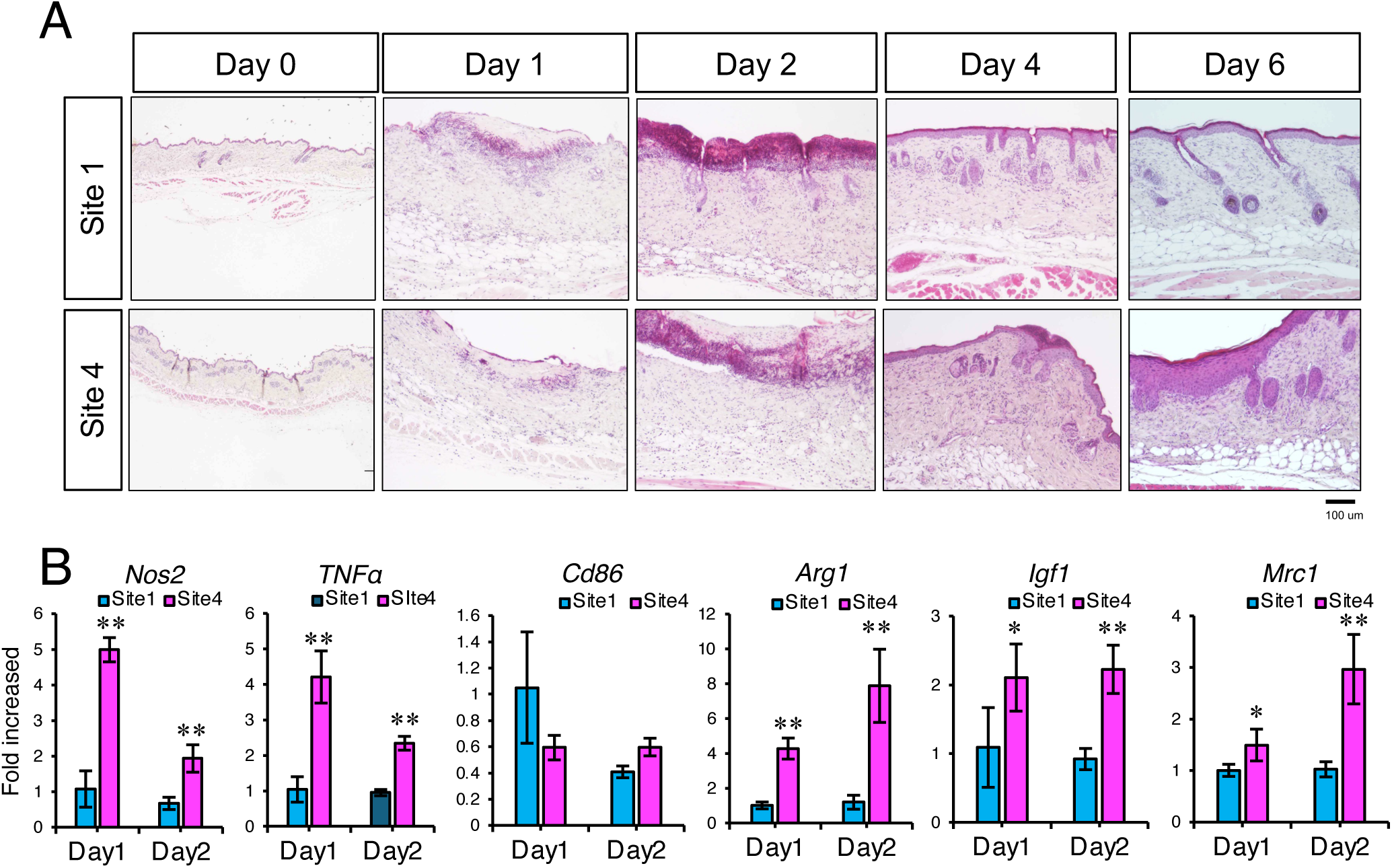
Histological and molecular responses of cranial and caudal dorsal skin during adhesive material-induced hair regrowth. Adhesive material was applied to Site 1 and Site 4, and histological changes and gene expression during the hair-regrowth process were analyzed. (A) Representative hematoxylin and eosin-stained images of Site 1 and Site 4 on Day 0, 1, 2, 4, and 6 after adhesive material application. Scale bar: 100 μm. n = 6. All images show representative examples. (B) Expression analysis of the inflammation-related genes *Nos2*, *Tnfα*, and *Cd86*, and the tissue-repair-related genes *Arg1*, *Igf1*, and *Mrc1*, on Day 1 and Day 2 after adhesive material application. Data are presented as mean ± SD. **P < 0.01 vs. Site 1 at the same time point (unpaired two-tailed Student’s t-test). n = 5 animals; qPCR was performed in technical triplicate.

Analysis of inflammation- and tissue-repair-related genes on Day 1 and Day 2 showed that the inflammation-related genes *Nos2*, *Tnfα*, and *Cd86* showed different expression patterns between Site 1 and Site 4. The tissue-repair- and macrophage-polarization-related genes *Arg1*, *Igf1*, and *Mrc1* also showed expression that differed by site and time point (Figure 5B). Together, these results indicate that Site 1 and Site 4 differ not only in baseline gene expression before adhesive material application but also in the subsequent inflammatory and tissue-repair response.

## Discussion

In this study, we found that the adhesive material-induced hair-regrowth response in mouse dorsal skin differs along the cranial-to-caudal axis. Even within the dorsal skin of a single animal, the telogen-to-anagen transition was induced earlier in the cranial region, whereas this response was delayed in the caudal region. This time difference was not limited to the onset of hair regrowth but was maintained through late anagen and the transition to catagen. In contrast, wound stimulation by full-thickness skin excision induced hair regrowth more slowly than adhesive material application, with variable presence, absence, and area of hair regrowth. Furthermore, cranial and caudal dorsal skin differed not only in gene-expression profile before adhesive material application but also in the early tissue changes and the expression of inflammation- and repair-related genes after application. Together, these results indicate that mouse dorsal skin is not uniform with respect to the hair-induction response, tissue response, and gene-expression profile, but instead shows regional differences along the cranial-to-caudal axis.

Yu et al. reported that Hoxc-dependent mesenchymal niche heterogeneity of dermal papilla cells alters the regenerative activity of hair follicle epithelial stem cells and contributes to site-specific hair-growth capacity [22]. In the present study, however, Hox gene expression was higher in the caudal region, where the hair-regrowth response was delayed, rather than in the cranial region, where the adhesive material-induced hair-regrowth response was early (Figure 3). Thus, at least in this experimental system, the cranial-to-caudal difference in hair-regrowth response cannot be simply explained by the expression pattern of the Hox genes examined.

Wnt/β-catenin signaling has been reported to be important for hair follicle stem cell activation, the telogen-to-anagen transition, and hair-cycle progression [2,10]. In this study, expression of the Wnt-signaling inhibitor *Sfrp4* was higher in the caudal region. Reduced Wnt signaling associated with increased *Sfrp4* expression in the caudal region may therefore be one factor contributing to the delayed hair-regrowth response in the caudal, compared with the cranial, region.

In addition, dermal and subcutaneous adipose tissue are known to be involved in hair-cycle regulation. Festa et al. showed that dermal adipocyte-lineage cells fluctuate with the hair cycle and contribute to the hair follicle stem cell niche [3]. In this study, expression of *Pparγ* and *Fabp4*, which reflect adipocyte differentiation and the presence of adipose tissue, was reduced in the caudal region (Figure 4E). This suggests that support for hair follicle activation from the adipose/mesenchymal niche may be relatively reduced in the caudal region.

Nevertheless, although hair regrowth was induced earlier in the cranial than in the caudal region, the cranial region also transitioned to catagen correspondingly earlier (Figure 1E). That is, rather than a large difference in the duration of anagen itself between the cranial and caudal regions, it is more likely that the timing of onset of the telogen-to-anagen transition differs between the two regions. In our previous study, adhesive-material-induced hair regrowth was suggested to arise through a wound-like inflammatory stimulus [8]. The cranial-to-caudal difference in the timing of hair-regrowth onset observed in this study may therefore reflect a difference in the reactivity of the early post-application inflammatory and tissue-repair response — which can act as a trigger for hair regrowth — rather than a difference in the intrinsic anagen-maintenance capacity of the hair follicles themselves. Indeed, in this study, the early tissue response and the expression of inflammation- and repair-related genes after adhesive application differed between the cranial and caudal regions. The underlying molecular mechanism, however, has not been identified and remains a subject for future investigation.

In the full-thickness skin excision model, the telogen-to-anagen transition after wound stimulation was slow, and the appearance of hair regrowth was unstable. In contrast, in the adhesive material model, nearly complete hair regrowth was obtained within a short period, consistent with the site of application. Small full-thickness skin excision wounds have been reported as a system that induces the telogen-to-anagen transition of resting hair follicle stem cells [6]; however, our results suggest that, compared with this existing full-thickness excision-based hair-induction system, the adhesive material model allows the telogen-to-anagen transition to be analyzed over a shorter period and with higher reproducibility, making it a useful system for hair-cycle research.

Both models are considered experimental systems in which wound stimulation activates resting hair follicles and induces their transition to anagen. However, why adhesive-induced hair regrowth is so much more reliable and extensive than that induced by full-thickness excision is an important question. In full-thickness excision, inflammatory stimuli spread from the surrounding wound margin, whereas in adhesive-induced hair regrowth, the stimulus is applied directly from the skin surface. Furthermore, because epidermal stripping occurs after adhesive application, the upper structure of existing hair follicles is also directly damaged, which may induce a subsequent repair and reconstruction process. Indeed, hair follicles have been reported to retain reconstructive capacity even when partially damaged: after plucking, sites of hair follicle epithelial damage are re-epithelialized and the follicle structure is reconstructed [5]. In addition, cell populations capable of regeneration may exist even in transected hair follicle fragments [17].

Taken together, these findings suggest that adhesive-induced hair regrowth may not simply arise from inflammatory stimuli spreading from the surrounding wound margin, but rather that epidermal stripping and damage to the upper hair follicle structure trigger a repair and reconstruction process in existing follicles, which in turn strongly promotes the telogen-to-anagen transition. Whether adhesive-induced hair regrowth primarily depends on epidermal stripping, damage to the upper follicle structure, the inflammatory response, or the follicle-reconstruction process itself will require further investigation. Clarifying these mechanisms may lead to the development of new hair-growth-promoting methods that efficiently induce the transition of resting hair follicles to anagen.

A limitation of this study is that the RNA-sequencing analysis comparing Site 1 and Site 4 (Figure 4) was performed using paired samples from only 2 animals (n = 2 animals total, with both Site 1 and Site 4 sampled from each). Although this paired within-animal design controls for inter-animal variability, the small number of animals limits the statistical power of the differential-expression analysis, and the set of differentially expressed genes reported here should be regarded as candidate region-specific pathways rather than a definitive transcriptomic signature; independent validation in a larger cohort, for example by qPCR in additional animals, will be needed to confirm these findings.

In conclusion, this study showed that the adhesive material-induced hair-regrowth response differs along the cranial-to-caudal axis of mouse dorsal skin. Dorsal skin is widely used in many skin-related research fields, including wound healing, dermatitis, skin barrier function, evaluation of topical agents, and hair growth/hair-cycle research. However, our results indicate that dorsal skin is not a uniform experimental region, but instead shows tissue responsiveness that differs according to anatomical position. In particular, in experimental systems that analyze the telogen-to-anagen transition of resting hair follicles using wound stimulation or adhesive application, regional differences within the dorsal skin may influence the interpretation of results. Therefore, in hair-regeneration and wound-healing research, dorsal skin should not be treated as a single uniform region; experimental design and interpretation of results should take cranial-to-caudal positional information into account. This study demonstrates the importance of considering regional differences within mouse dorsal skin in hair-regeneration research, and shows that the adhesive material-induced hair-regrowth model is a useful experimental system for analyzing local hair-cycle responses with high reproducibility.

